# Functional relevance of dual olfactory bulb in olfactory coding

**DOI:** 10.1101/588996

**Authors:** Praveen Kuruppath, Li Bai, Leonardo Belluscio

## Abstract

Bilateral convergence of external stimuli is a common feature of vertebrate sensory systems. This convergence of inputs from the bilateral receptive fields allows higher order sensory perception, such as depth perception in the vertebrate visual system and stimulus localization in the auditory system. The functional role of such bilateral convergence in the olfactory system is mostly unknown. To test whether each olfactory bulb contributes a separate piece of olfactory information, and whether information from the bilateral olfactory bulb is integrated, we synchronized the activation of olfactory bulbs with blue light in mice expressing channelrhodopsin in the olfactory sensory neurons and behaviorally assessed the relevance of dual olfactory bulb in olfactory perception. Our findings suggest that each olfactory bulb contributes separate components of olfactory information and mice integrate the olfactory information from each olfactory bulb to identify an olfactory stimulus.

**Significance statement:** Identifying an odor is the first step in olfactory coding, as it is critical for the survival of most animals. Previous studies have shown that bilateral olfactory bulbs help rodents to localize the odor source and navigate accordingly. But It is still unclear whether the bilateral olfactory information plays any role in determining odor identity. Here for the first time, using optogenetics and behavioral experiments, we demonstrate that each olfactory bulb provides distinct olfactory information, and rodents integrate information from the two bulbs to identify an odor.

## Introduction

The perception of odor is crucial for the survival of most animals because it is essential for navigation, finding food sources and avoiding predators. Convergence of sensory inputs from a bilateral receptive field is a fundamental aspect of the biological sensory systems to extract information about the environment. In the visual, auditory and somatosensory system, this bilateral convergence is well established for depth perception (Anzai et al., 2011; Ohzawa et al., 1997), sound localization (King et al., 2001; Konishi, 2003) and object localization (Shuler et al., 2001, 2002). The significance of such bilateral convergence and the functional relevance of dual olfactory bulb in the olfactory system remain unclear.

Odorant identity and the concentration are two fundamental aspects of olfactory information, representing the quality and the quantity of the odorant signal. Animals move towards the increasing odorant concentrations to locate food and mates and avoid predators by moving away following decreasing odorant concentrations. To achieve this effect, animals may rely on a comparison of odorant information sampled through the bilateral symmetric nostrils (Esquivelzeta Rabell et al., 2017; Raman et al., 2008). Previous studies have shown that rodents integrate bilateral cues from the nostrils to localize the odorants sources (Catania, 2013; Rajan et al., 2006). However, it is still unknown whether bilateral olfactory inputs are integrated and perceived as a single olfactory information.

Here, we precisely controlled the sensory inputs to the olfactory bulbs (OB) with blue light in tetO-ChIEF-Citrine mice line, where channelrhodopsin-2 is expressed in all the olfactory sensory neurons (OSNs) and studied the behavioral responses to the light pulse stimulation. We found that each olfactory bulb provides separate olfactory information and the final identity of the olfactory stimulus is the composite of dual olfactory bulb.

## Materials and methods

### Experimental animals

All animal procedures conformed to National Institutes of Health guidelines and were approved by the National Institute of Neurological Disorders and Stroke Institutional Animal Care and Use Committee. Mice were bred in-house and were maintained on a 12 h light/dark cycle with food and water ad libitum.

The tetO-ChIEF-Citrine line, was generated from pCAGGS-I-oChIEF-mCitrine-I-WPRE (7.7kb; Roger Tsien, UCSD), which contains the coding sequence for mammalian-optimized ChIEF fused to the yellow fluorescent protein Citrine, at the National Institute of Mental Health Transgenic Core Facility (Bethesda, MD) as previously described (Cheetham et al., 2016; Lin et al., 2009). The OMP-tTA knock-in mouse line expressing the tetracycline transactivator protein (TTA) under the control of the OMP-promoter was a generous gift from Dr. Joseph Gagos. Experimental animals were OMP-tTA+/− / tetO-ChIEF-Citrine+/− (OMP-ChIEF), generated by crossing heterozygous tetO-ChIEF-Citrine (tetO-ChIEF-Citrine+/−) with homozygous OMP-tTA mice (OMP-tTA-/-).

### Genotyping

OMP-ChIEF pups were identified by the visualization of fluorescence in the nose and OB of P0-P2 pups under epifluorescence illumination.

### Animal preparations

Data were collected from 12 OMP-ChIEF mice and 4 wild type mice. Experimental animals were prepared as described previously (Sparta et al., 2012; Ung and Arenkiel, 2012). Briefly, mice were anesthetized with an intraperitoneal injection of Ketamine/Xylazine mixture 100 and 10 mg/kg body weight, respectively. Each animal was fixed with a stereotactic frame with the head held in place by a bar tie to each temporal side of the skull. The animals were kept warm with hand warmers (Grabber, Grand Rapids, MI, USA). Surgery was started when the animal showed no movement in response to foot pinching. A craniotomy was performed above the skull over each OB. Fiber optic pins were implanted over each OB as described previously (Sparta et al., 2012; Ung and Arenkiel, 2012). Mice were injected with Ketoprophen (5mg/kg) immediately after the surgery. Animals were allowed to recover in their home cage for 1 week.

### Behavioral procedure and training

#### Light stimulation and Foot shock avoidance training

Behavioral training began after the animals recovered from the surgery (1 week). Training was performed on a modified ‘Y’ maze with two equally sized open arms and one permanently closed arm. Each open arm was independently paved with electric grid shock floor. The mice were connected to a 400 μm core-diameter optical fiber attached to a 473 nm solid-state variable-power laser (LaserGlow Technologies, Toronto, Canada, Figure. 1A) and allowed to habituate in the ‘Y’ maze for 15 minutes. The time spent in each arm of the ‘Y’ maze was recorded and assessed for any particular arm preference. The preferred arm for the mice was selected as the light-zone and the opposite arm served as the safe-zone. If the mice did not have any preference to any particular arm, the light zone was randomly selected. After the habituation, mice were re-introduced to the ‘Y’ maze for the 10 minutes training session. A reinforcement training was performed every day prior to the test session to enhance the learning. The training consists of light stimulation followed by a mild foot shock. The light stimulation and the foot shock were delivered in the light zone when the mice completely entered into that zone (Figure 1B). The mice have free access to the safe zone to escape from the foot shock. Light stimulation, consisting of a train of 10 light pulses of 50 ms duration with an interval of 150 ms, was externally triggered by a Master-8 timer (A.M.P.I, Jerusalem, Israel). The output power of the light pulses was measured and adjusted to 20-22 mW. The mild foot shock (0.65mAmps, 5 s) generated by a stand-alone shock generator (Med Associates, USA) was delivered 2 s after the light stimulation by Master-8 timer. The mice were trained to move to the safe zone when the light stimuli and the foot shock were delivered in the light zone.

**Figure 1.**
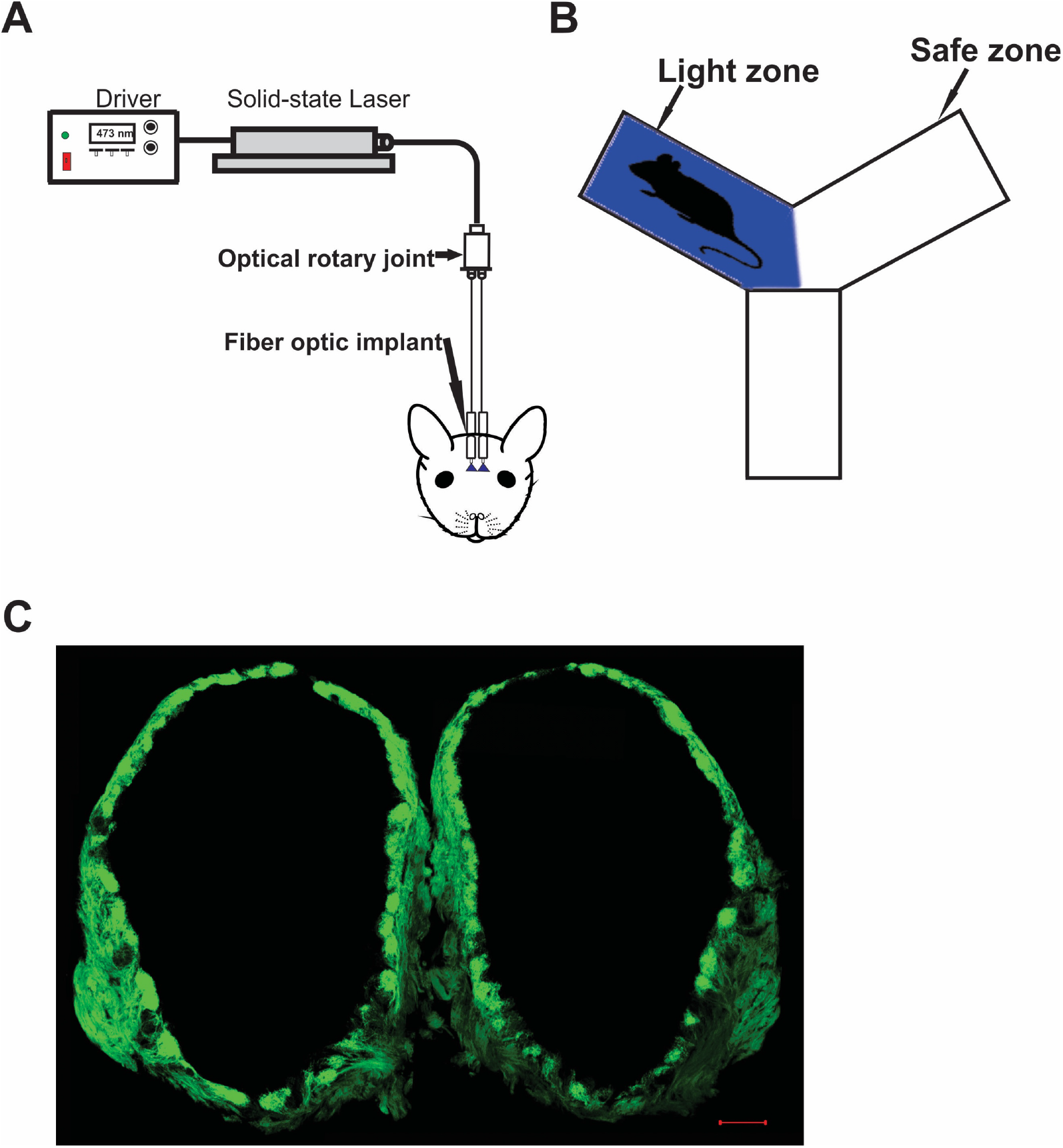
Optogenetic stimulation of Olfactory bulb. **A**, Schematic diagram of the functional system. **B**, Behavioral setup. Light zone indicates the area where the light stimulation is delivered. Safe zone indicates the area to which the mice can escape from the light stimulation and foot shock. **C**, Fluorescence image of the olfactory bulb in an OMP-ChIEF mouse. Green indicate the expression of ChR2 tagged with a green fluorescent protein (ChR2-GFP). Scale bar: 100 μm.

#### Light zone avoidance test

The testing began 60 minutes after the behavioral training. Before the testing, the electric grid shock floor was removed from the ‘Y’ maze, so the mice did not experience any foot shock during the test session. The activity of the mice in the ‘Y’ maze was evaluated in blocks of 3 trials and each trial lasted 15 minutes. In the first trial, the mice were allowed to explore the arena without any light stimulation and assessed the baseline behavior. The time spent in each arm was calculated to evaluate any particular arm preferences after the foot shock training session. For the following trials, light zone and safe zone were selected as mentioned previously. The time spent in each arm was calculated by tracking the animal’s movement in the ‘Y’ maze using Any-maze video tracking software (Stoelting, IL, USA). A heatmap was also generated for each trial which shows the amount of time mice spent in different parts of the arena. A range of colors indicate the total time spent in the area with blue as the shortest and the red as the longest time.

#### Perfusion and immunohistochemistry

At the end of behavioral testing, the mice were deeply anesthetized with 200 mg/kg ketamine and 20 mg/kg xylazine and transcardially perfused with ice-cold PBS followed by 4% paraformaldehyde (PFA). OB were dissected out and embedded in 10% gelatin and post-fixed/cryopreserved in 15% sucrose and 2% PFA in PBS overnight at 4 °C, cryopreserved in 30% sucrose in PBS for 24 h at 4 °C and flash frozen in 2-methyl butane on dry ice. Coronal sections were cut using a cryostat (Leica Microsystems) at 50 μm and stored at −80 °C.

For immunohistochemistry, free-floating sections were incubated for 20 min in 1% sodium borohydride in TBS, blocked in the blocking medium containing 5% horse serum, 0.1% gelatin and 0.5% Triton-X100 for 1 h, and incubated with OMP primary antibody (Goat, Wako, 1:1000) in 3% horse serum and 0.2% Triton-X100 for 24 h at 4 °C, then Donkey-anti-goat Cy3 secondary antibody (Jackson ImmunoResearch, 1:600) for 2 h at room temperature. Sections were mounted in Vectashield mounting medium (Vector Laboratories, CA). Images were acquired using a Zeiss LSM 510 laser scanning confocal microscope.

#### Statistical analysis

All statistical analyses were done by Graph Pad prism software (Graph-pad, San Diego, CA). Statistics are displayed as mean ± SEM. Unpaired *t* test was used for the comparison. Differences were determined significant when p< 0.05.

## Results

### Olfactory information is distinct in each olfactory bulb

Previous studies have shown that odor information is projected unilaterally (Shepherd, 2004) in vertebrate olfactory system, indicating that odors may be encoded separately in each hemisphere. To investigate the unilateral coding of olfactory information, we optically stimulated ChR2 expressing OSNs in each olfactory bulb (Figure 1C) and assessed the behavior in response to light stimulation. Here, we performed foot shock avoidance training paired with either left or right OB stimulation. In our study, the optical fiber implanted on each OB were in different locations. So, the light stimulation on each OB activate different set of glomeruli, eliciting different olfactory information in each OB. After training, we tested the light zone avoidance response to ipsilateral and contralateral OB stimulation. First, we tested the mice’s baseline behavior and calculated the time spent in each arm by allowing the mice to freely explore the arena. The aim of the baseline behavior analysis was to check if the mice have any preference to a particular arm of the ‘Y’ maze after the foot shock training. Our baseline data show that mice spent almost equal amounts of time in both arms of the ‘Y’ maze (Left zone - 471.6 ± 25.57 s, Right zone-428.4 ± 25.57 s, Figure 2A). Then, we delivered the light stimulation to ipsilateral OB. We found that, mice avoided light zone during the ipsilateral OB stimulation and spent most of their time in the safe zone (Light zone - 74.67 ± 13,07 s, Safe zone - 825.3 ± 13,07 s, Figure 2B, 2C, Movie S1).

**Figure 2.**
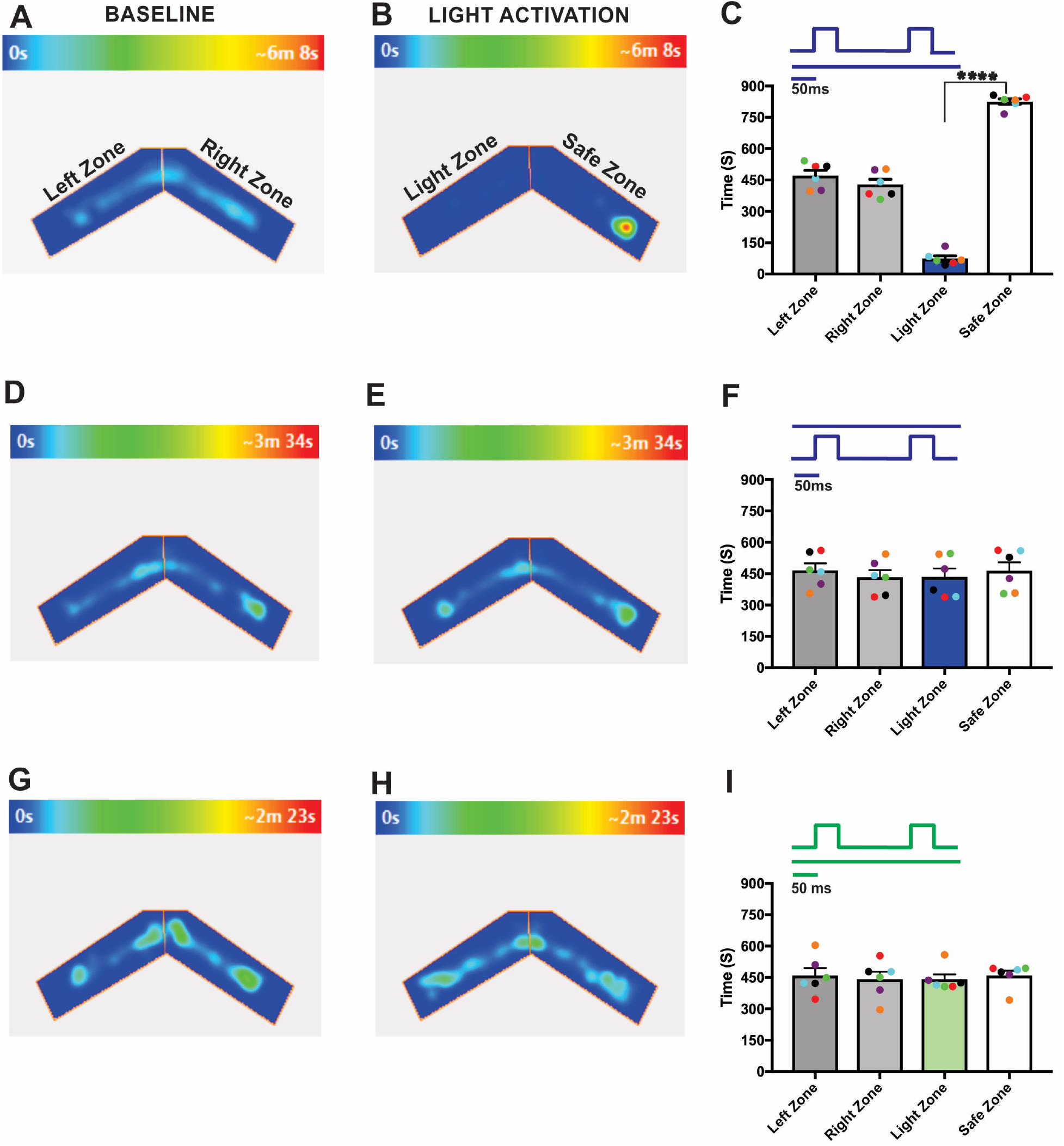
Distinct odor information from individual OB. **A**, **B**, an example of heat-map showing animal’s position in Y maze during baseline (**A**) and ipsilateral OB stimulation (**B**). **C**, Average amount of time explored in each zone in baseline and ipsilateral OB stimulation trials. **D**, **E**, Heat-map of animal’s position during baseline (**D**) and contralateral OB stimulation (**E**). **F**, Average amount of time spend in each zone in baseline and contralateral OB stimulation trials. **G**, **H**, Heat-map of animal’s position during baseline (**G**) and ipsilateral green light stimulation (**H**). **I**, Average amount of time spend in each zone in ipsilateral green light stimulation. (****P<0.0001, n=6 animals).

Next, we performed light stimulation on the contralateral OB. Here we found that the time spent in each arm during the baseline behavior trial (Left zone - 466.4 ± 33.25 s, Right zone - 433.6 ± 33.25 s, Figure 2D) and the light activation trial (Light zone - 435.3 ± 39.93 s, Safe zone - 464.7 ± 39.93 s, Figure 2E) were almost equal, indicating that the mice did not identify the foot-shock linked olfactory information. Thus, they did not avoid the light zone during contralateral OB stimulation (Figure 2F, Movie S2). This suggest that each OB provides two separate light-induced olfactory information which are encoded independently in each hemisphere and the mice are able to discriminate this distinct olfactory information.

To verify that the responses we observed from the light stimulation was the result of activation of the ChR2 expressing neurons and not from the use of light itself as a visual cue, we used green light as the light source, which does not activate ChR2. During the green light stimulation, we did not see any significant behavioral difference from the baseline behavior (Left zone - 459 ± 36.13 s, Right zone - 441 ± 36.13 s, Light zone - 441 ± 23.88s, Safe zone - 459 ± 23.88 s, Figure 2G-I), confirming that mice did not use any visual cues to perform the task. We also performed the test with wild-type mice (Figure 3A, B) and confirmed that visual cues are not involved in solving the behavioral task. Together, these results indicate that, OMP-ChIEF mice were using only the light activation of the olfactory system as the cue to solve the behavioral task rather than light detection via other modalities.

**Figure. 3.**
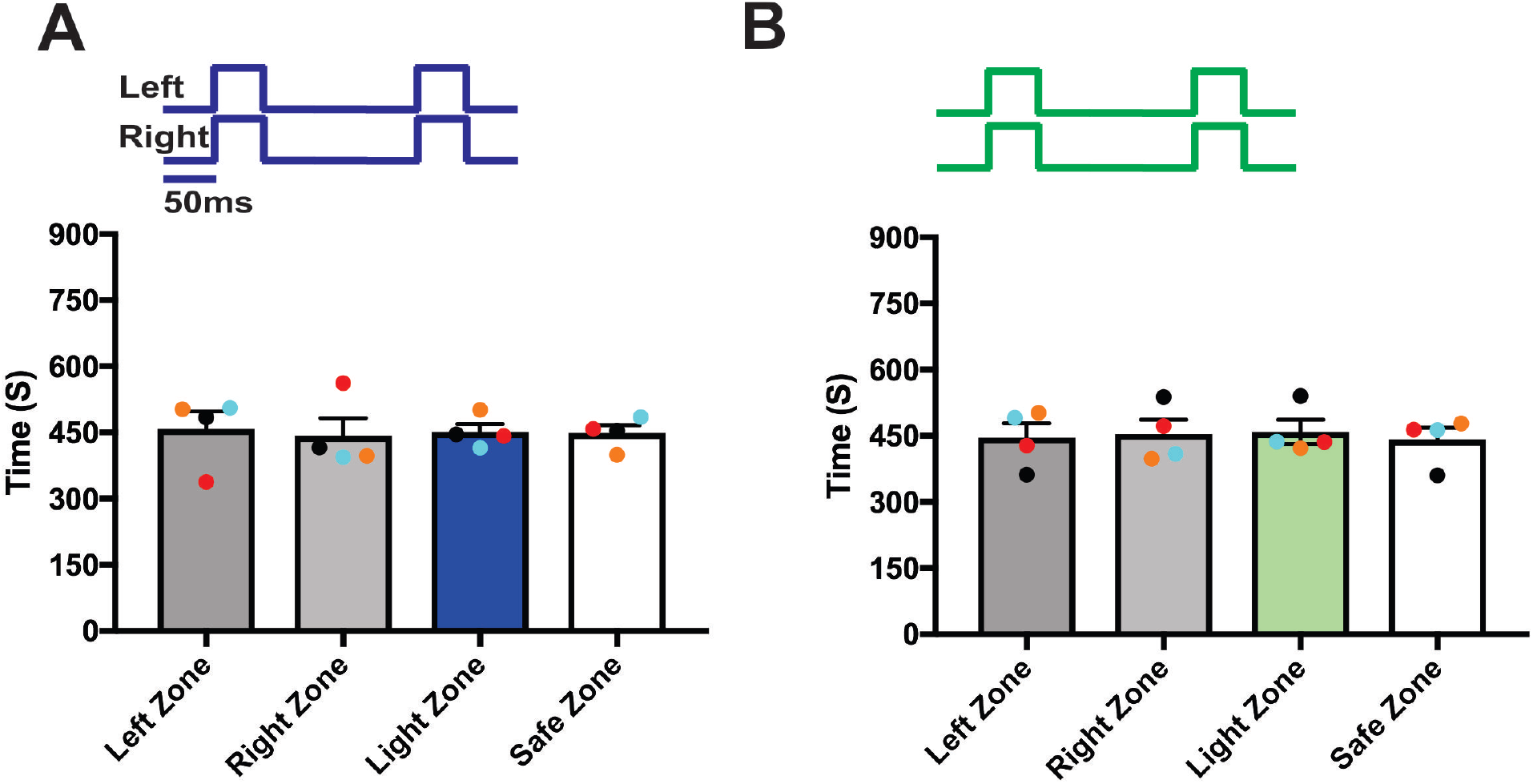
Light stimulation did not activate the olfactory system of the wild-type mice. **A**, Average amount of time spent in each zone in baseline and experiment trials during blue light stimulation. **B**, Average amount of time spent in each zone in baseline and experiment trials during green light stimulation.

### Bilateral integration of olfactory information

Previous tracer injection studies have shown that olfactory information is exchanged to the contralateral olfactory bulb through anterior olfactory nucleus pars externa (AONpE) (Yan et al., 2008). Yet, it is unknown whether the olfactory information from both olfactory bulbs is integrated in odor perception. To assess if the bilateral olfactory information is integrated in olfactory perception, we delivered synchronized light stimulation to both olfactory bulbs of a unilaterally trained mice and checked if the mice can identify the foot-shock associated olfactory information from the synchronized dual bulb stimulation. We found that during the baseline behavior trial, and during the synchronized dual OB stimulation trials, mice visited both arms equally (Left zone - 438.2 ± 20.67 s, Right zone - 461.8 ± 20.67 s, Light zone - 447.8 ± 39.57 s, Safe zone - 452.2 ± 39.57 s, Figure 4A, Movie S3). This indicates that mice failed to identify the olfactory information linked to the foot-shock and did not avoid the light zone, suggesting that mice integrate the information from the ipsilateral and contralateral olfactory bulbs during synchronized bilateral OB stimulation and perceive it differently from one bulb alone.

**Figure 4.**
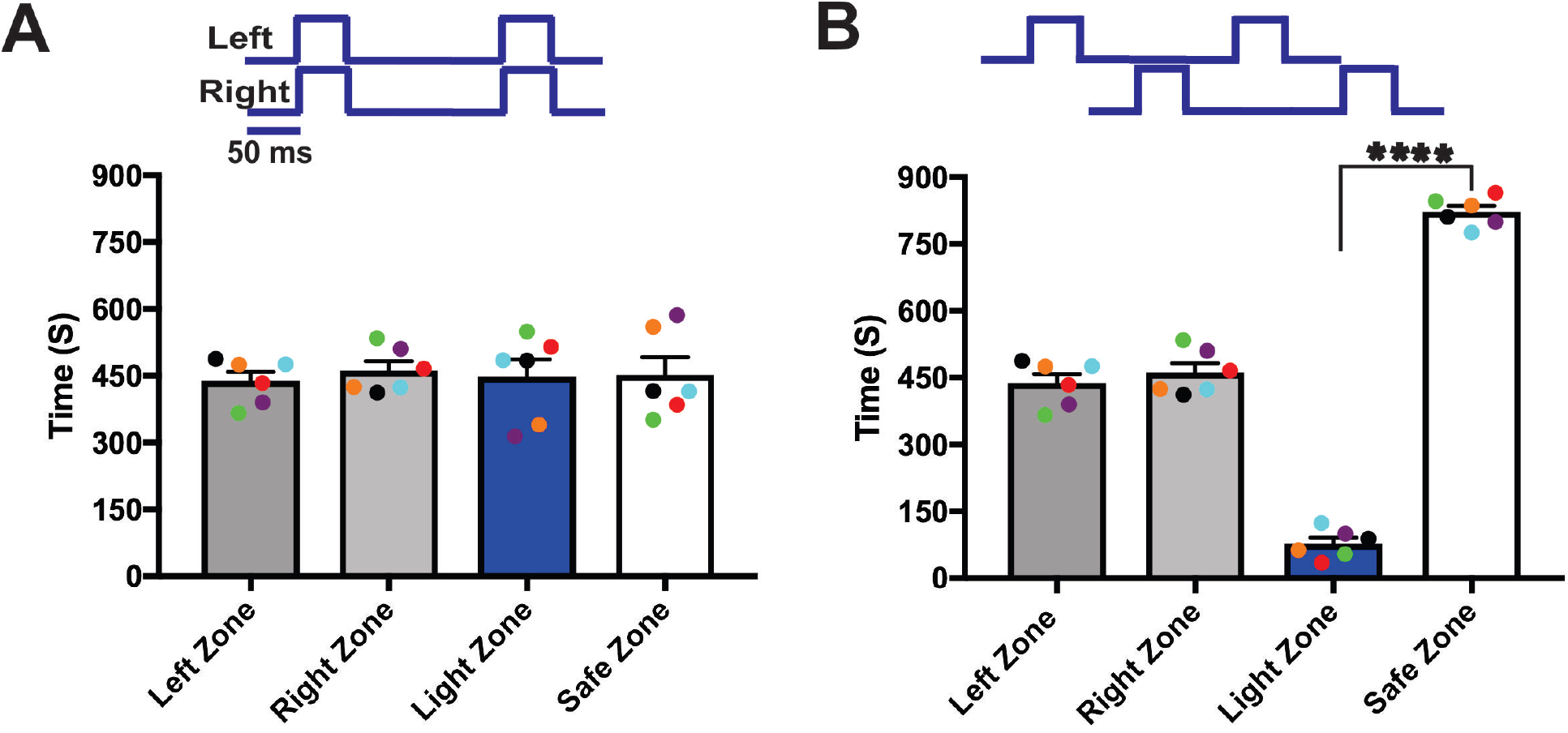
Synchronized bilateral OB stimulation alter the Unilateral odor identity. **A**, **B**, mean time explored during the synchronous and asynchronous OB stimulation to the unilaterally trained mice. (****P<0.0001, n=6 animals).

To verify the bilateral integration, we delivered the light stimulation asynchronously with a 50 ms delay between ipsilateral and contralateral OB stimulation. This assumed that asynchronous OB stimulation did not allow the mice to integrate the olfactory information because each olfactory bulb gets the light stimulation at a different time. We found that when stimulated asynchronously, mice identified the distinct light-driven olfactory information paired with the foot-shock and avoided the light zone (Light zone - 77.58 ± 13.36 s, Safe zone - 822.4 ± 13.36 s, Figure 4B, Movie S4), suggesting that during asynchronous stimulation olfactory information is processed separately. Thus, the mice can clearly identify the ipsilateral and contralateral olfactory information.

### Unified perception of bilateral olfactory inputs

Sensory information from the bilateral receptive fields is perceived as unified in most sensory system. However, the presence of such a unified perception in the vertebrate olfactory system is still unknown. To determine the unified perception of bilateral olfactory information, we performed light stimulation and foot-shock avoidance training on a new group of animals (n=6). Here we delivered light stimulation to both olfactory bulbs simultaneously and paired it with foot shock. After the training, we tested the response of mice in the light zone avoidance test during synchronized bilateral OB stimulation. Our results show that when both olfactory bulbs were synchronously stimulated mice avoided the light zone, indicating that mice can identify the olfactory information linked to the foot-shock (Light zone - 83.67 ± 8.891 s, Safe zone - 816.3 ± 8.891 s Figure 5A, Movie S5). Then we tested if the mice can identify the foot-shock associated olfactory information when we deliver the light stimulation asynchronously. This assumes that if the mice were relying on the olfactory information from either left or right olfactory bulb to avoid the light zone, mice could identify the foot-shock linked olfactory information and avoid the light zone. In contrast to this prediction, we found that mice failed to identify the olfactory information linked with foot-shock and continued to stay in the light zone (Light zone - 450.2 ± 42 s, Safe zone - 449.8 ± 42 s, Figure 5B, Movie S6). Our results suggest that mice integrate the olfactory information from both olfactory bulbs during synchronized stimulation and perceive it as a single olfactory information, but during the asynchronized stimulation, olfactory information from each olfactory bulb was processed separately and perceived it as two independent pieces of information.

**Figure 5.**
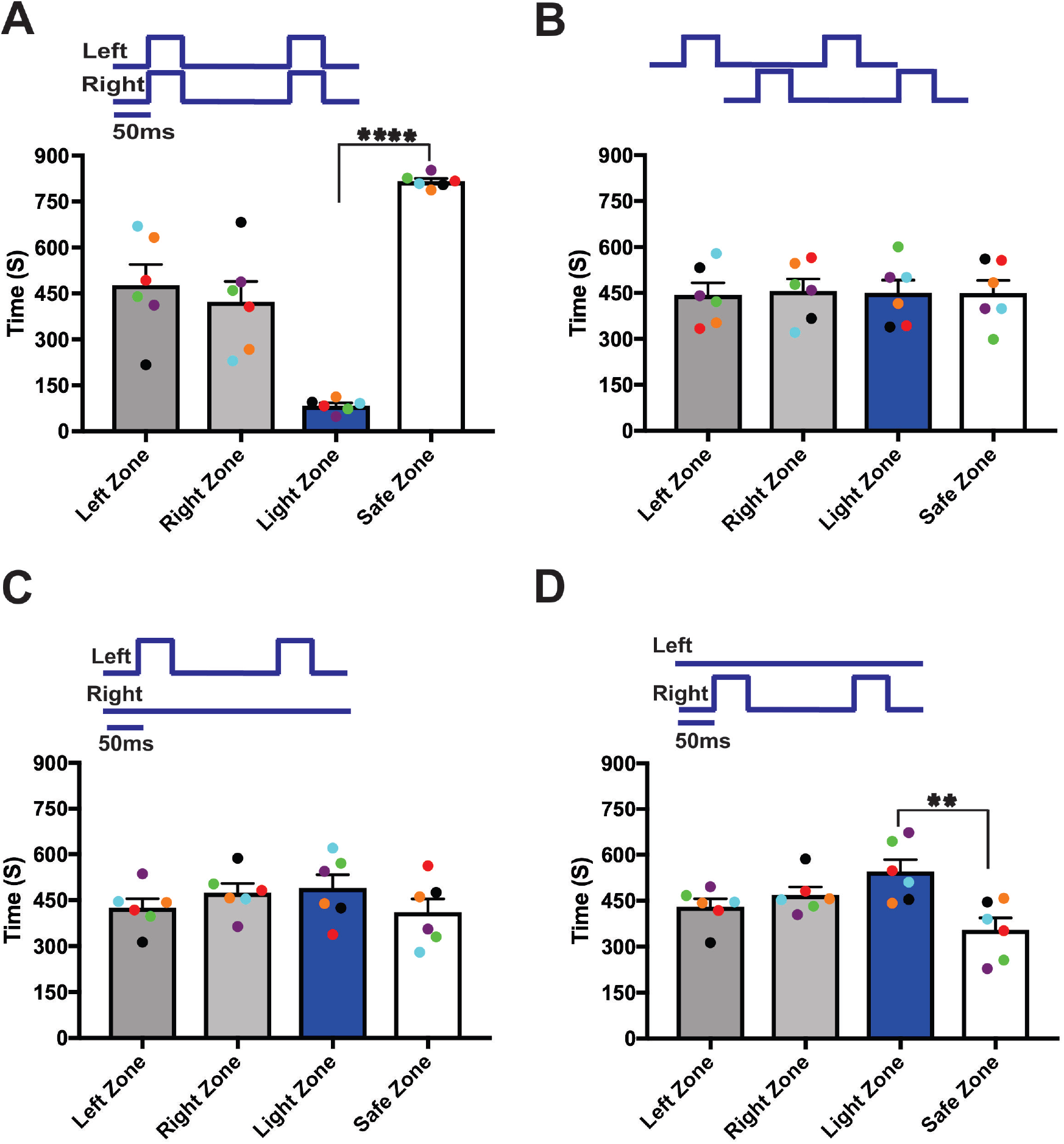
Unified perception of synchronized bilateral olfactory inputs. **A**, **B**, mean time explored during the synchronized and asynchronized bilateral OB stimulation to the bilaterally trained mice. **C**, **D**, mean time explored during unilateral light stimulation. (**P<0.01, ****P<0.0001, n=6 animals).

We then tested whether the mice can identify the foot-shock linked bilateral olfactory information from single olfactory bulb stimulation. Here we independently stimulated left and right olfactory bulb of the bilaterally trained mice. We found that during the independent activation of each olfactory bulb, mice did not avoid the light zone, but spent equal or more time in the light zone as in baseline behavior trials (Figure 5C, D). This suggests that unilateral stimulation of each olfactory bulb provides only a piece of bilateral olfactory information. This result confirms our previous assumption that each OB provides two separate pieces of information, which are encoded independently in each hemisphere.

Finally, we compared the OMP positive OSNs in 12wk old OMP-ChIEF control, experimental and wild type mice and examined whether the light stimulation induced any reduction in the ChR2 expression and the degeneration of OMP positive neurons in the wild type and experimental mice. Our immunohistochemistry results show that light stimulation did not induce any reduction in the ChR2 expression or degeneration of the OMP positive OSNs (Figure 6A – C).

**Figure 6.**
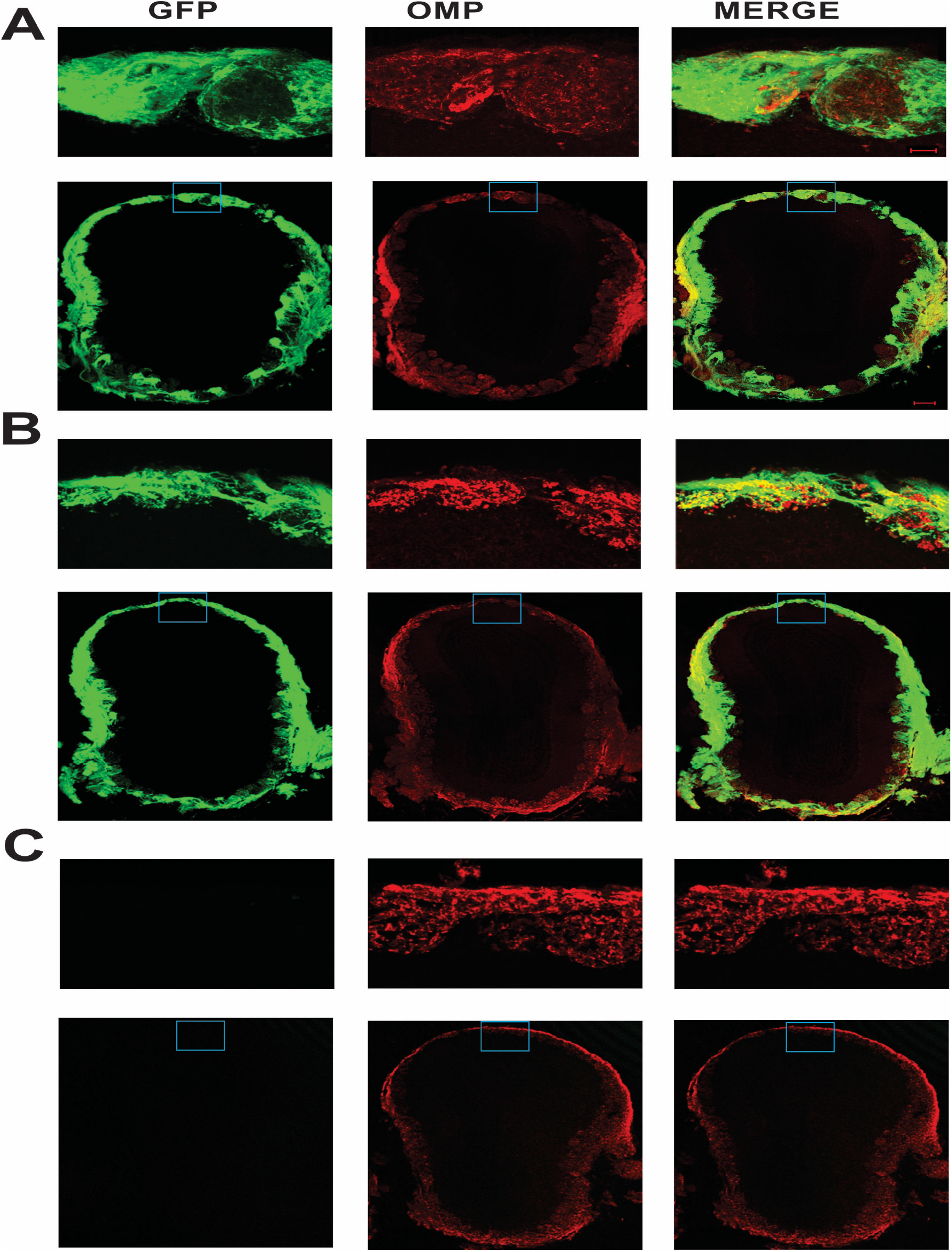
ChR2 expression in OMP positive neurons. **A**, ChR2 expression in OMP positive OSNs in 12wk old OMP-ChIEF control animal. **B**, ChR2 expression in OMP positive OSNs in 12wk old OMP-ChIEF experimental animal. **C**, no ChR2 expression on the OMP positive neurons of 12wk old Wild-type animal. Scale bars: **A-C**, 100 μm; inset, 20 μm.

Together, our results show for the first time that each OB provide distinct olfactory information, which can be combined and perceived as a single odor stimulus.

## Discussion

Several studies have shown that rodents use bilateral cues to localize odor source for navigation, finding food sources and avoiding danger. While these studies report the significance of bilateral sampling for accurate odor source localization (Catania, 2013; Esquivelzeta Rabell et al., 2017; Louis et al., 2008; Rajan et al., 2006), none of the studies reported the significance of bilateral integration of olfactory information for odor identity. Here for the first time, we report that rodents combine the synchronously sampled bilateral olfactory information for odor identity. Previous studies have reported that olfactory information is projected unilaterally (Shepherd, 2004), suggesting that odorant information are encoded separately in each hemisphere, thus providing unique olfactory information in each olfactory bulb. Consistent with this, our study also provides evidence for distinct olfactory information processing in each hemisphere. We show that when stimulated independently, olfactory information in ipsilateral and contralateral olfactory bulb is unique and the mice can clearly identify the olfactory information from the two bulbs (Figure 2C, 2F).

What is the functional relevance of the dual olfactory bulbs in olfactory information processing is still an open question. A recent study in Drosophila found that olfactory receptor neurons (ORNs) project bilaterally to both sides of the brain. When an odor activates the antennal lobe asymmetrically, ipsilateral central neurons begin to spike a few milliseconds earlier at a higher rate than the contralateral neurons, allowing the fly to localize the odor direction by comparing the bilateral response differences (Gaudry et al., 2013).

Previous studies in rodents also reported that isofunctional glomeruli are interconnected not only within the hemisphere but also between the hemispheres through AON, creating a mirror symmetric olfactory map in each hemisphere (Grobman et al., 2018; Mainland et al., 2002; Mombaerts et al., 1996; Ressler et al., 1994; Schoenfeld and Macrides, 1984; Uchida et al., 2000; Wilson, 1997). Together, these reports suggest that dual olfactory bulb enhance the odor source localization and navigation. However, Grobman and co-workers (Grobman et al., 2018) suggest that bilateral projection is not necessary to get a bilateral response difference. Instead, a unilateral projection can result in a strong bilateral response difference. They show that a unilateral projection pathway generates a higher bilateral response difference than the bilateral pathway, and the odor localization is easier in the unilateral projection pathway than the bilateral pathway. Therefore, they suggest that the main role of bilateral projection is not odor localization but the sharing of odor identity information across the hemispheres.

In agreement with their reports, we propose that light stimulation of a single OB forms a neural representation of an odor’s molecular identity in both hemispheres. In addition to that, we also propose that the final odor identity is represented not only by the combinatorial activation of specific ORs as previously reported (Malnic et al., 1999; Saito et al., 2009), but also by the integration of the olfactory information from the contralateral olfactory bulb.

Based on the previous studies and our own results, we suggest that one of the main roles of dual olfactory bulb is the coding of odor identity. Since we did not investigate the role of dual olfactory bulbs for odor localization, we do not know from our study whether the dual olfactory bulbs contribute any valuable information about the source of an odor. Dual olfactory bulbs may enable odor localization through the bilateral projection. But one important thing that should be noted here is that, if the ipsilateral and contralateral odor responses are of same magnitude, bilateral response differences will be same. In such cases odor localization would be impossible. Further research is required to better understand how olfactory cortical neurons integrate the dual olfactory information for odor identity and odor localization.

In their natural environments, animals confront more complex problems such as, having to identify the quality, quantity and the complexity of an odor mixture. The use of suitable physiological and psychophysical paradigms will be a crucial step for further understanding the complexity of neural coding in the olfactory system.

