## Supplemental Movie Legends for "Functional relevance of dual olfactory bulb in olfactory coding"

**Movie S1.** Unilateral foot shock trained mice avoid light zone during ipsilateral olfactory bulb Stimulation.

**Movie S2.** Unilateral foot shock trained mice did not avoid light zone during contralateral olfactory bulb Stimulation.

**Movie S3.** Unilateral foot shock trained mice did not avoid light zone during synchronized bilateral olfactory bulb stimulation.

**Movie S4.** Unilateral foot shock trained mice avoid light zone during asynchronous bilateral olfactory bulb stimulation.

**Movie S5.** Bilateral foot shock trained mice avoid light zone during synchronized bilateral olfactory bulb stimulation.

**Movie S6.** Bilateral foot shock trained mice did not avoid light zone during asynchronous bilateral olfactory bulb stimulation.
